# Alzheimer’s disease-relevant tau modifications selectively impact neurodegeneration and mitophagy in a novel *C. elegans* single-copy transgenic model

**DOI:** 10.1101/2020.02.12.946004

**Authors:** Sanjib Guha, Sarah Fischer, Gail VW Johnson, Keith Nehrke

**Affiliations:** University of Rochester, Department of Anesthesiology & Perioperative Medicine, Rochester, NY; University of Rochester, Department of Medicine, Nephrology Division, Rochester, NY

**Keywords:** Alzheimer’s disease, *C. elegans*, tau, neurodegeneration, post-translational modifications

## Abstract

**Background:** A defining pathological hallmark of the progressive neurodegenerative disorder Alzheimer’s disease (AD) is the accumulation of misfolded tau with abnormal post-translational modifications (PTMs). These include phosphorylation at Threonine 231 (T231) and acetylation at Lysine 274 (K274) and at Lysine 281 (K281). Although tau is recognized to play a central role in pathogenesis of AD, the precise mechanisms by which these abnormal PTMs contribute to the neural toxicity of tau is unclear.

**Methods:** Human 0N4R tau (wild type) was expressed in touch receptor neurons of the genetic model organism *C. elegans* through single-copy gene insertion. Defined mutations were then introduced into the single-copy tau transgene through CRISPR-Cas9 genome editing. These mutations included T231E and T231A, to mimic phosphorylation and phospho-ablation of a commonly observed pathological epitope, respectively, and K274/281Q, to mimic disease-associated lysine acetylation. Stereotypical touch response assays were used to assess behavioral defects in the transgenic strains as a function of age, and genetically-encoded fluorescent biosensors were used to measure the morphological dynamics and turnover of touch neuron mitochondria.

**Results:** Unlike existing tau overexpression models, *C. elegans* single-copy expression of tau did not elicit overt pathological phenotypes at baseline. However, strains expressing disease associated PTM-mimetics (T231E and K274/281Q) exhibited reduced touch sensation and morphological abnormalities that increased with age. In addition, the PTM-mimetic mutants lacked the ability to engage mitophagy in response to mitochondrial stress.

**Conclusions:** Limiting the expression of tau results in a genetic model where pathological modifications and age result in evolving phenotypes, which may more closely resemble the normal progression of AD. The finding that disease-associated PTMs suppress compensatory responses to mitochondrial stress provides a new perspective into the pathogenic mechanisms underlying AD.

## BACKGROUND

Alzheimer’s disease (AD) is the most common degenerative brain disease in the aged population. It is characterized by the progressive decline of cognition and memory, as well as changes in behavior and personality (1). One of the key pathological hallmarks of AD is neurofibrillary tangles (NFTs), which are primarily composed of abnormally modified tau (2). Tau isolated from AD brain exhibits a number of posttranslational modifications (PTMs); including increases in phosphorylation and acetylation at specific residues (3, 4). While it is clear that tau is central to AD pathogenesis, the concept of large insoluble NFTs in AD and other tauopathies being the principle mediators of neuronal toxicity has been gradually abandoned (5). Instead, toxicity appears to result from soluble or oligomeric forms of tau that exhibit increased, disease-associated phosphorylation and acetylation at specific residues altering its turnover and function (6, 7).

Studies to date have provided evidence that phosphorylation of tau at Threonine 231 (T231) occurs early in the evolution of tau pathology; for example, increased staining for this epitope is observed in “pre-tangle” neurons (8). Further, increased phospho-T231 tau was observed in neurons differentiated from iPSCs of sporadic AD cases (9). Phosphorylation of tau at specific sites causes significant changes in tau structure (10, 11) and impairs microtubule binding (12, 13). In addition, phosphorylation of tau at T231 precedes the formation of tau oligomers (7, 14), which likely contribute to tau toxicity (15).

As with phosphorylation, abnormal tau acetylation also likely plays a critical role in tauopathies (16–18). There are data indicating that acetylation inhibits binding of tau to microtubules, enhances tau accumulation by preventing degradation and promotes the aggregation of tau in neurons (19–21). In particular, increased expression of tau acetylated at K274 and K281 appears to result in mislocalization of tau, destabilization of the cytoskeleton in the axon initial segment, and synaptic dysfunction (20, 22). Altogether, these experiments suggest a potential role for tau acetylated at K274/281 in AD pathogenesis. While these studies indicate that modifications of human tau at specific residues play a pivotal role in mediating tau dysfunction, the precise mechanism by which specific tau PTMs contribute to the toxicity of soluble tau forms is still unclear.

Mitochondrial dysfunction is a characteristic of many neurodegenerative diseases including AD (23, 24), and over expression of human full length tau or mutant human tau contributes to mitochondrial dysfunction in AD animal models (25, 26). Mitochondria play a complex role in the cell - they not only generate most of the energy needed to support the various neuronal functions (27), but also are mediators of homeostatic processes that are necessary for neuronal health (28). Although it is likely that tau pathology affects mitochondrial biology, the underlying mechanisms are not well understood, nor it is known how tau modified at disease relevant sites differ from its wild type form in causing mitochondrial abnormalities leading to neurodegeneration.

To understand the role of tau in the context of AD *per se*, tau transgenic models have been developed in *C. elegans* (29–31), *D. melanogaster* (32, 33) and mice (34, 35) by overexpression of human wild-type full-length tau (25), tau with mutations that result in frontotemporal lobar degeneration (FTLD) (35), or a tau with a risk factor mutation for AD FTLD such as A152T (31). Studies utilizing these transgenic animals have made important contributions to the field, but over-expression of tau can potentially lead to synthetic toxic or gain-of-function phenotypes, and this caveat must always be kept in mind when extrapolating results to the human disease.

Here, we attempted to circumvent the limitations associated with tau overexpression by taking advantage of single-copy genome insertion methodology in the genetic model organism *C. elegans* (36). Using this methodology, human tau was expressed in a defined set of mechanosensory neurons that mediate a stereotypical behavioral output (37). To interrogate the effects of pathologic PTMs in this system, CRISPR-Cas9 gene editing was used to introduce AD-associated phosphorylation mimicking (T→E) or a non-phosphorylatable (T→A) mutation at the T231 position of the wild-type tau isoform, or alternatively acetylation mimicking (K→Q) mutations at the K274 and K281 positions of the wild type tau isoform. A combination of behavioral assays and fluorescent biosensors were used to study the impact of tau and mutant tau expression on neuronal morphology and mitochondrial phenotypes, with the advantage of being able to assess age-dependence in a relatively short time frame (38, 39).

Our results clearly demonstrate that wild-type tau has little effect at baseline, but that AD-relevant tau PTMs selectively impact sensory neuron function and morphology and mitochondrial handling. Moreover, age exacerbates defects in one of the tau mutant strains, but not the others. This leads us to conclude that using our single copy tau model confers the ability to discern between the pathological consequences of individual tau mutants with unprecedented precision. Surprisingly, AD-associated tau mutants also completely suppressed paraquat-induced mitophagy, supporting the idea that pathological modifications of tau results in dysfunctional responses to stress, including perhaps the stress of aging.

## METHODS

### Plasmid construction

Briefly, pBJ1 codes for the fluorescent photo-convertible protein Dendra2 (40), cloned downstream of the *mec-7* promoter in a pFH6.II *C. elegans* expression vector (41). pBJ2 adds the coding sequence for tau (0N4R) inserted downstream and in-frame with Dendra2. pBJ5 and pBJ6 are derivatives of pBJ1 and pBJ2, respectively, with the tau expression cassette sub-cloned into pCFJ151 (Addgene) to generate MosSCI inserts at the ttTi5605 loci in *C. elegans* chromosome II (42). pSKG1 contains a *mec-4* promoter driving the expression of *C. elegans* codon-optimized mito-mKeima (courtesy of Dr. C. Rongo, Rutgers University).

### *C. elegans* strain generation

The wild-type background strain was Bristol-N2. Other strains used here include, KWN177 *rnyIs14* [*Pmec-4::mCherry*], KWN796 *rnyEx336 [pSKG1 (Pmec-4::mito-mKeima), pCFJ90 (Pmyo-2::wCherry), pCI (pha-1+)*]. Transgenic strains for single copy gene expression were generated using MosSCI insertion (42) into ttTi5605 on Chromosome II via established protocols (36), and include the following: KWN169, *rnySi26 [Pmec-7::Dendra2; unc-119+] II*; KWN167, *rnySi24 [Pmec-7::Dendra2::Tau-T4; unc-119+] II*. Both strains were sequenced completely through the insertion site and were outcrossed at least four times to the lab N2-Bristol stock. CRISPR-Cas9 gene editing was used to introduce site-specific mutations into the rnySi24 tau coding region via a co-CRISPR strategy and oligonucleotide-mediated HDR using purified Cas9 RNP injection (43, 44). Targeting crRNAs were from Dharmacon and were complexed to scaffolding RNAs for Cas9, with genomic recognition sites as follows:

Tau T231, 5’ACGGCGACTTGGGTGGAGTA3’;

Tau K274/281, 5’GCACCAGCCGGGAGGCGGGA3’.

Single stranded oligonucleotide directed repair templates were:

Tau T231A ssODN,

5’GTCCCTTCCAACCCCACCCACCCGGGAGCCCAAGAAGGTGGCCGTGGTCAGAGCCCCACCCAAGTCGCCGTCTTCCGCCAAGAGCCGCCTGCAGA3’

Tau T231E ssODN,

5’GTCCCTTCCAACCCCACCCACCCGGGAGCCCAAGAAGGTGGCCGTGGTCAGAGAGCCACCCAAGTCGCCGTCTTCCGCCAAGAGCCGCCTGCAGA3’

Tau K274/281Q ssODN,

5’CGGCTCCACTGAGAACCTGAAGCACCAGCCGGGAGGCGGGCAAGTGCAGATAATTAATAAGCAGCTGGATCTTAGCAACGTCCAGTCCAAGTGTGGCTCAAAGGATA3’

In all cases, HDR would be predicted to disrupt the PAM, but leave the coding sequence potential outside of the desired amino acid substitution intact. Repair at T231 also disrupted a BtsaI site, while repair at K274/281 created a new PvuII site. These modifications could be detected via restriction analysis of genomic PCR products and were used to screen *dpy-10* co-CRISPR mutants for edits with primers:

Tau geno-F1, 5’-AAAGACACCACCCAGCTCTG-3’

Tau geno-R1, 5’TGTTGCCTAATGAGCCACAC3’,

Following isolation of homozygous tau mutants, editing was confirmed via genomic PCR sequencing, and the mutants were crossed out of the co-CRISPR’d *dpy-10* mutant background. The final strains are KWN788 *rnySi51 [Tau-T4 (T231A) *rnySi24] II*, KWN789 *rnySi52 [Tau-T4 (T231E) *rnySi24] II*, KWN790 *rnySi53 [Tau-T4 (K274Q; K281Q) *rnySi24] II*. For crossing tau MosSCI strains into various genetic backgrounds, Dendra2 fluorescent was used to guide selection of homozygous mutants, and PCR genotyping was used to confirm homozygosity with primers specific to the ttTi5605 loci, including:

MosSCI ttTi5605-F, 5’GTTTTTGATTGCGTGCGTTA3’

MosSCI ttTi5605-R, 5’ACATGCTTCGTGCAAAACAG3’

MosSCI ttTi5605 insert-F, 5’CATCCCGGTTTCTGTCAAAT3’

Other strains included KWN791 *rnySi51 II*; *rnyIs14,* KWN797 *rnySi26 II*; *rnyIs14* KWN798 *rnySi24 II, rnyIs14,* KWN800 *rnySi52 II*; *rnyIs14,* KWN801 *rnySi53* II; *rnyIs14* KWN802 *rnySi26 II*; *rnyEx336* KWN803 *rnySi24 II*; *rnyEx336*, KWN804 *rnySi52 II*; *rnyEx336*, KWN805 *rnySi53 II*; *rnyEx336*, KWN806 *rnySi51 II*; *rnyEx336*.

### *C. elegans* strains growth and maintenance

Nematodes were maintained at 20°C on Nematode Growth Media (NGM) plates made with Bacto Agar (BD Biosciences). The plates were seeded with live *E. coli* OP50-1 bacterial strain (cultured overnight at 37°C at 220 rpm) and allowed to grow overnight, as previously described (45). For experimental assays, after synchronization by standard procedure with sodium hypochlorite, 4^th^ larval stage (L4) hermaphrodites (characterized by the appearance of a “Christmas tree vulva”) were selected and moved to test plates. The day after moving was considered adult day 1, and animals were assayed on day 3 and day 10. Animals were transferred daily to avoid mixed population until they stop laying eggs.

### Blinding of experiments and replicates

Insofar as possible, experimentalists were blinded to genotype. Data in the figures generally represents the pooled results of three experimental replicates with either two technical replicates per condition or two independent researchers blindly analyzing the data, with the total number of animals or neurons scored reported as N, as indicated.

### Locomotory Rate Assay

Assay plates were prepared using standard procedures (46). Synchronized day 3 and day 10 adult animals were assayed for the actual experiment. For well-fed animals, locomotory rate was measured by removing 5 animals from original plate and transferring them to an assay plate. Five minutes after transfer, the number of body bends in 20 secs intervals was sequentially counted for each of the 5 animals on the assay plate and then repeated the same thing for next set of animals in a different assay plate.

### Thrashing Assay

A drop of 2% agarose (ultraPURE® agarose) was poured over the glass slide and allowed to dry and then 20μl of M9 was poured on it. Age-synchronized animals were picked to that drop of M9 buffer. After 2 min in M9, thrashing rates were assessed via videography on a stereo dissecting scope. A single thrash was defined as a complete change in the direction of the body down the midline. Animals that were motionless for 10 secs were discarded from the analysis (47).

### Touch sensitivity assay

The behavioral response to being touched by an eyelash was adapted from an assay previously described (48, 49). The animals were touched anteriorly specifically behind the terminal bulb of the pharynx with the eyelash, 10 times per animal, with a 10 sec gap between each touch. Typically, if the animal demonstrates an omega turn or if it reversed its direction after an anterior touch, the animal was scored as giving positive response. Touch response percentage was generated by the amount of times an animal responded to the touch stimulus over the total number of times they were touched.

### Life span analysis

After alkaline hypochlorite treatment, synchronized L1 animals were placed on freshly grown OP50-1 seeded NGM plates. 15 animals from the 4^th^ larval stage (L4) were transferred to a small (35mm) individual seeded NGM plate with a total 3 plates for each genotype. Each day they were transferred to new plates to avoid mixing of populations until they stopped producing offspring. Simultaneously the worms were counted alive visually or with gentle prodding on the head. Animals were censored in the event of internal hatching of larva, body rupture or crawling off the plate. The experiment was conducted at 20°C temperature and scored until all the worms died (50).

### Mitochondrial stress assays

For paraquat (PQT) mild stress assays, synchronized 2-day old adult and 9-day old adult hermaphrodites were exposed to 8 mM PQT (51, 52) in NGM plate for overnight at 20°C. Animals were picked from the respective (treated and control) plates the next day and imaged, as described below.

### Neurodegeneration assay

For imaging, animals were mounted by placing them in 3% agarose pads on glass slides and immobilized with 1 mM tetramisole hydrochloride (Sigma). Imaging was performed using Confocal Laser Scanning Confocal microscope (Olympus 1X61) and FV10-ASW 4.1 software. All images were acquired under the same exposure conditions with a 20x objective, and for each experimental replicate, all genotypes were represented and imaged that day. In analysis of touch neurons, *Pmec-4:: mCherry* expressing animals scored positive for the presence of extra neuronal processes when a visible mCherry-labeled branch was observed emanating from the posterior portion of ALM cell body. Similarly, ALM / PLM neuron pairs were scored as overextended when the PLM neurite extended anterior to the ALM cell body (53, 54). Other defects in axonal morphology were assigned to one of the following classes of neuronal abnormality: broken or gap in the axon structure, blebbed or bead like structure on the axon body, misguided or wavy shaped axon (55).

### Mitophagy assay

A strain containing mito-mKeima (56, 57) expressed specifically in touch cells was used for assay. Animals were mounted on 2% agarose pads on glass slides and immobilized with 1 mM tetramisole hydrochloride before imaging. Imaging was performed using a Nikon Eclipse inverted microscope coupled to a six channel LED light source (Intelligent Imaging Innovation, Denver, CO), an ORCA-Flash4.0 V2 Digital CMOS camera (Hamamatsu Photonics, Bridgewater Township, NJ) and Slidebook6 software (Intelligent Imaging Innovation, Denver, CO). All images were acquired under the same exposure conditions and each experiment was imaged in one session. The PLM cell body was identified by their position toward the posterior of the animal, near the tail and was focused with a 100x oil immersion lens under visible light using DIC contrast. 600-nm+ emissions were captured first following excitation at 550-nm and then immediately thereafter at 440-nm, keeping light intensity and exposure times constant between images. Images were quantified using Slidebook6 software by selecting the ROI, measuring the mean intensities for both channels and subtracting the background intensity. Mitophagy index was obtained by calculating the dual excitation 550-nm/440-nm ratio.

### Mitochondrial Morphology assay

Mitochondrial morphology was analyzed using images acquired on a florescence microscopy rig as described above, from animals expressing mito-mKeima in the mechanosensory touch cells, acquired at 440-nm excitation, 600 nm+ emission. The morphological features were categorized into four distinct groups: 1) a network of long interconnected mitochondria with tubular-reticular or normal morphology, 2) visible fragmentation of the network, but lacking aggregates, 3) fragmentation consisting of short round mitochondria, but no more than one visible aggregate, 4) short round mitochondria comprising the majority of the population, with more than one large aggregate (58). Two investigators independently analyzed subsets of images and compared results to ensure the reproducibility of the analysis.

### Statistical Analysis

All statistical analyses were conducted using Prism 8.0 (GraphPad Software), with alpha-error level of *p < 0.05* considered to be significant. Data were averaged and represented as mean ± standard error (mean ± SEM) unless otherwise noted. In general, group differences were analyzed with either one-way or two-way ANOVA depending upon the variables. Fisher’s exact test was used to obtain p-values for the categorical data on pathologic neuronal morphology. Differences in lifespan were assessed by Mantel-Cox log rank analysis, and mitochondrial morphology data, which was categorical with four levels, was assessed using a Wilcoxen signed rank test. The sample sizes were based on those found previously in the laboratory to provide appropriate power for discerning phenotypic differences among genotypes. Graphs were plotted in Prism 8.0 (GraphPad Software) and Microsoft Excel.

## RESULTS

### Single-copy tau mutants that mimic AD-associated PTMs impact behavior

Tau expression via conventional extrachromosomal transgenic arrays in *C. elegans* has been shown to severely impact neuronal morphology and function (30). Here, in an attempt to circumvent potential caveats related to overexpression, novel transgenic AD models were engineered using single-copy Mos-transposon mediated insertion of a tau expression cassette into the worm genome (36, 42). The *mec-7* promoter was used to drive the expression of 0N4R tau (59, 60) as a translational fusion with the fluorescent protein Dendra2 (40) in mechanosensory touch neurons ALML/ALMR, AVM, PLML/PLMR, and PVM (Fig. 1E-H), which mediate the behavioral response to light touch. The 0N4R fusion to Dendra2 will be referred to hereafter as TauT4. Dendra2 was also expressed alone (Fig. 1A-D), and these negative control strains responded to light touch similarly to the wild-type N2 strain at both day 3 and day 10 of adulthood (Fig. S1).

**Figure 1.**
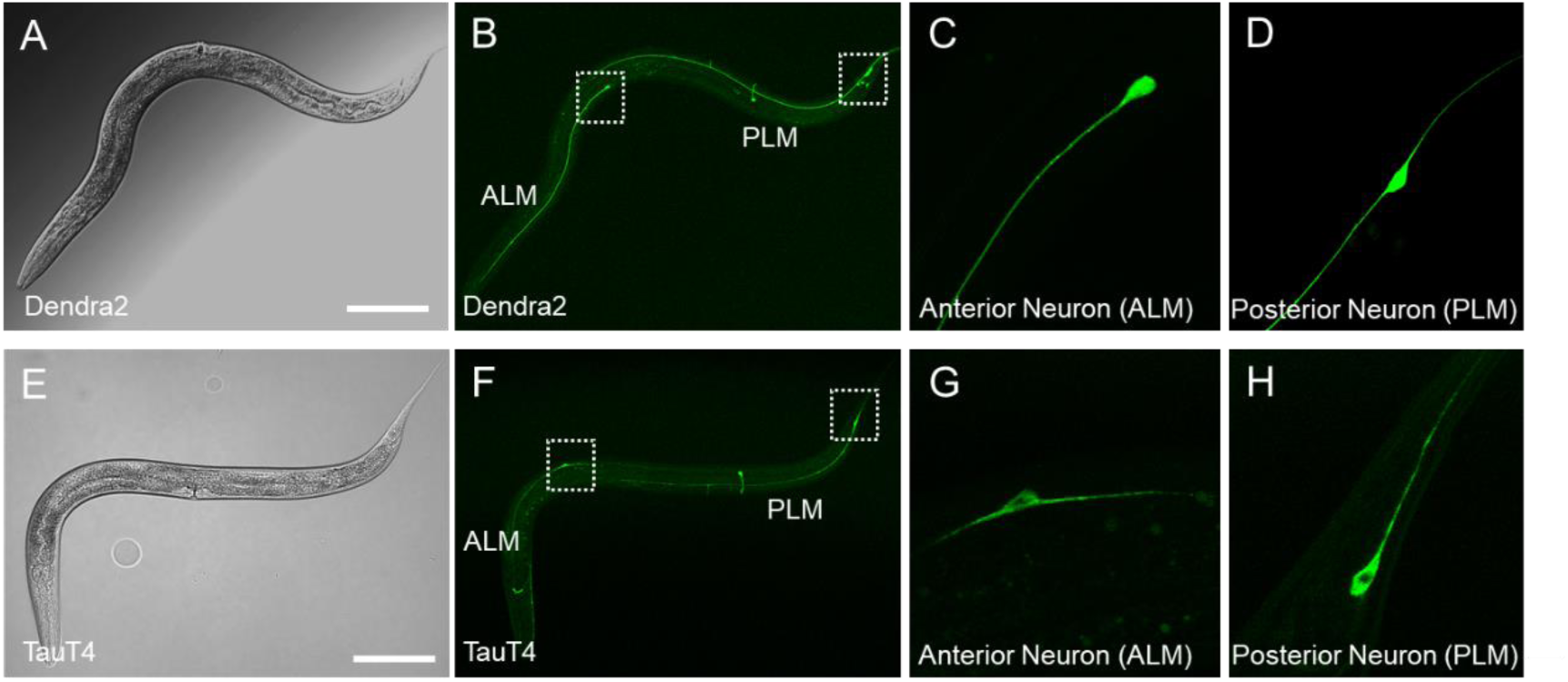
Expression of Dendra2 and TauT4 from a single-copy transgene in *C. elegans* touch neurons. DIC (A, E) and confocal fluorescent images (B, C, D, F, G, and H) are shown of L4 larval worms expressing single-copy transgenes coding for Dendra2 (B, C, D) or a Dendra2::TauT4 translational fusion (F, G, H). The transgenes are driven by the *mec-7* promoter in ALM(L/R) and PLM(L/R) neuron pairs, and also in AVM and PVM neurons, which are not considered further here. Panels C, G and D, H are magnifications of ALM and PLM respectively in the areas encompassed by the white boxes in panels B and F. Scale bars: 25 µm. ALM is <u>A</u>nterior <u>L</u>ateral <u>M</u>icrotubule and PLM is <u>P</u>osterior <u>L</u>ateral <u>M</u>icrotubule cells, mechanosensory neurons that mediate behavioral responses to light touch to the body wall within the receptive fields defined by their projections.

Surprisingly, TauT4 worms exhibited normal touch responsiveness as both young day 3 post-reproductive adults (Fig. 2B) and older day 10 adults (Fig. 2C). In order to address the effect of tau PTMs, CRISPR-Cas9 gene editing (43, 44) was used to introduce phosphomimetic T231E, phosphoablation T231A, and acetylmimetic K274/281Q mutations into the TauT4 ORF (Fig. 2A). For simplicity, these mutants will be referred to as T231E, T231A and K274/281Q. Our results clearly demonstrate that T231E exhibited subtle but significant defects in touch responsiveness at both day 3 and day 10, while K274/281Q was different from the Dendra2 control only at day 10 (Fig. 2B,C). However, between day 3 and day 10, the touch sensitive phenotype of K274/281Q worsened significantly *(p = 0.01)*. This may indicate either a ceiling effect of T231E or a sensitized K274/281Q progression with age. In contrast, T231A was indistinguishable from TauT4. The differences between the disease-associated mutants and TauT4 represents a novel observation and a first-in-kind platform for studying the effect of pathologic tau modifications in the absence of baseline defects. Finally, since survival plots of the various strains used in this work were statistically indistinguishable, we were able to rule out any phenotypic age-dependence being due to a change in lifespan (Fig. S2E, F).

**Figure 2.**
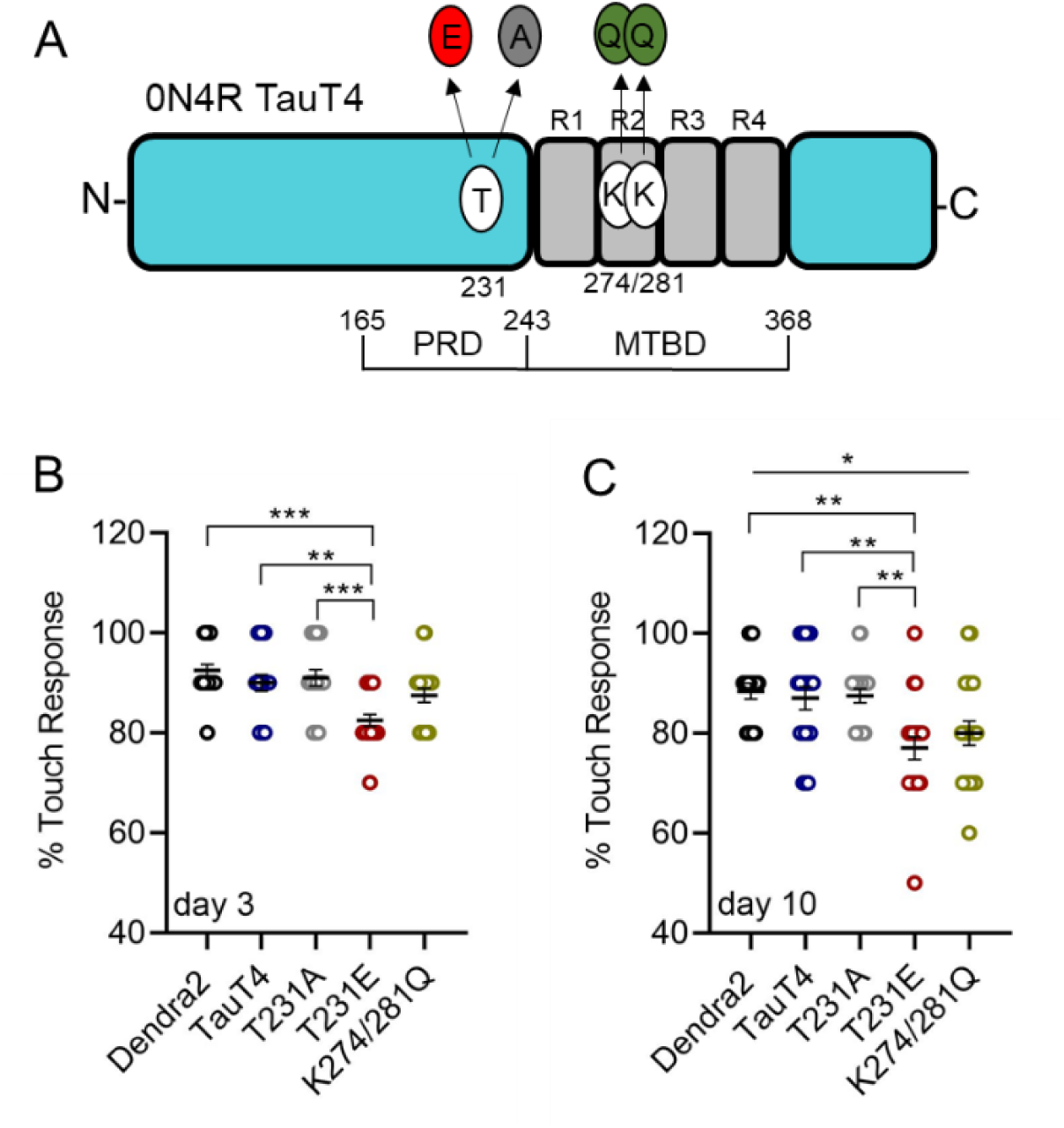
Tau mutations mimicking posttranslational modifications to T231 and K274/281 impact touch sensitivity in a single-copy transgenic *C. elegans* model. A) Schematic of TauT4 protein, with the proline-rich domain (PRD), microtubule-binding domain (MTBD), and repeats R1-R4 denoted, along with individual amino acids that were mutated by CRISPR/Cas9 editing. The numbering scheme is based upon Tau-441, the longest of the alternatively spliced human brain isoforms, as is the convention in the field (i.e. T231 is not the 231^st^ amino acid in the 0N4R tau variant, which lacks two N-terminal domains, but is instead positioned at amino acid 173). Touch sensitivity was quantified by measuring responsiveness to light touch in transgenic Dendra2, TauT4, T231A, T231E and K274/281Q mutant strains at day 3 (B) and at day 10 (C) of adulthood (d0 is when the worms enter their reproductive phase). Data were calculated as percent responsiveness following ten repetitive light touches to the anterior body, and are plotted with the mean ± SEM. Statistical analysis was by one-way ANOVA followed by Tukey’s multiple-comparisons test (N = 20 animals), with \**P*<0.05, \*\**P*<0.01 and \*\*\**P*<0.001 denoting significance between bracketed samples. Each circular point represents a value obtained from a single animal – note that many of the points overlap.

We also evaluated several other stereotypical behavioral measures that have been shown to be influenced by age but do not involve touch cell neurons, including thrashing in liquid (Fig. S2A, B) and basal locomotion on solid media (Fig. S2C, D). Taken together, these data suggest that the effect of pathological, AD-relevant tau expression in touch sensory neurons is restricted to the behavioral response to light touch.

### T231E and K274/281Q mutants cause age-dependent abnormalities in neurite morphology

Normally, touch neurons are organized into precise anterior and posterior receptive fields, defined by the physical architecture of sensory neurites from ALM(L/R) and PLM(L/R); these neurites extend along the anterior or posterior half of the body, respectively, but do not overlap (54). Aging phenotypes in touch receptor neurons include a low incidence of morphologic defects, such as increased neurite overlap due to an overextension defect (55). We investigated whether single-copy expression of Dendra2, TauT4 or the PTM mutants exacerbated these defects. A transgene consisting of an integrated *Pmec-4::mCherry* expression cassette was used to visualize the touch neurons using confocal microscopy (Fig. 3A-F). We found that T231E strongly and significantly increased the incidence of overextension from ~4% to ~40% by day 3 of adulthood (Fig. 3G). However, the TauT4 and K274/281Q mutants were not significantly different from Dendra2 controls in day 3 adults (Fig. 3G).

**Figure 3.**
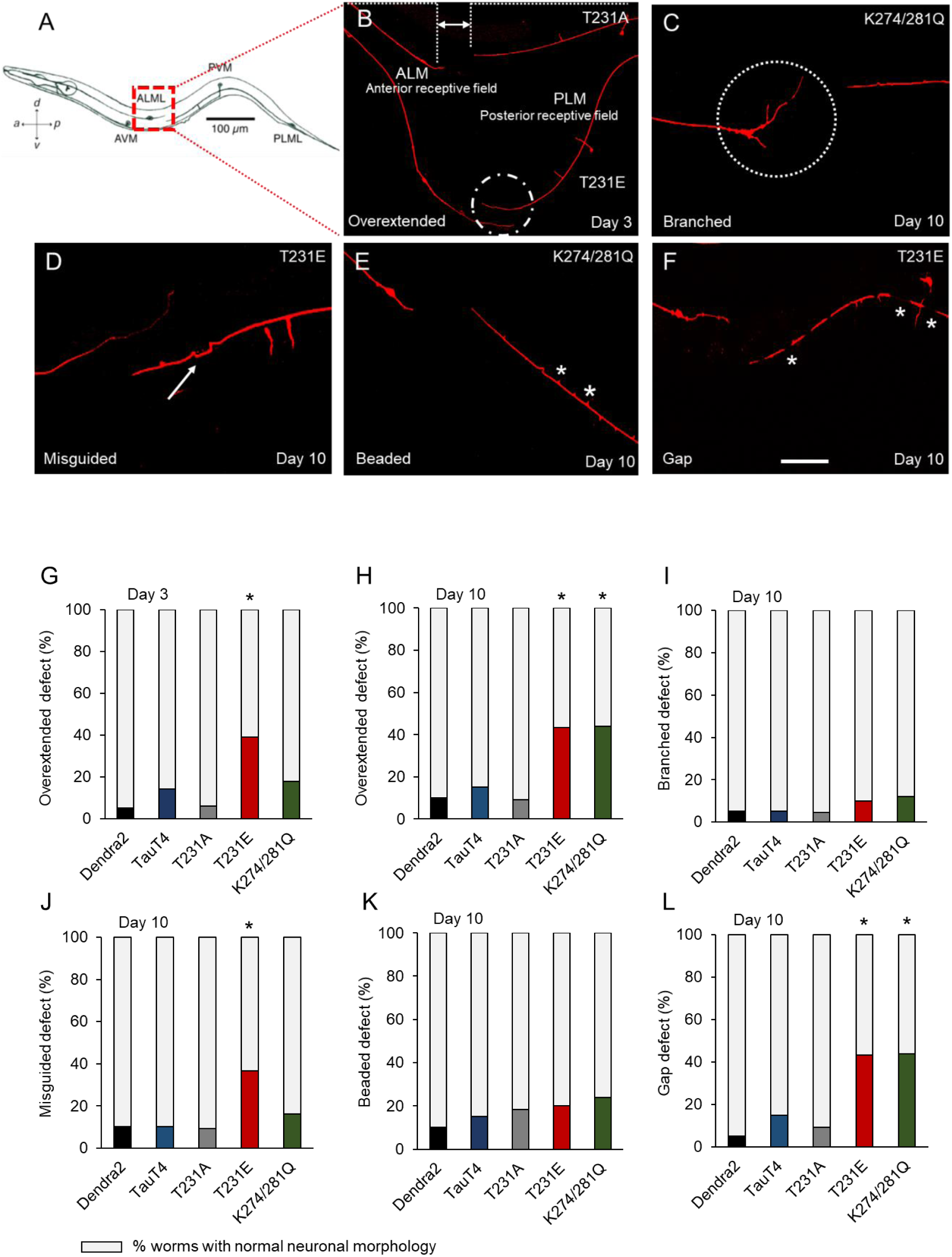
Abnormal touch receptor neurite morphology in the PTM mutants. (A) Schematic of a hermaphrodite animal. Mechanosensory neurons pairs ALM(R/L) and PLM(R/L) are present on both left and right sides of the animal, but only one of each pair is depicted. In wild type animals, neurites projecting from ALM and PLM do not overlap with each other, but instead divide the animal’s body into two distinct receptive fields, as indicated (modified from 79). (B) Neurons were visualized using a *Pmec-4::mCherry* fluorescent reporter. Two animals lying side-by-side are shown here. The top animal is from the phospho-null strain (T231A) and the bottom animal is from the phospho mimetic strain (T231E). The normal separation between the ALM and PLM neurites, represented by the area between the dashed lines in T231A, is replaced by overlapping neurites in T231E, as demarcated by a dashed circle. (C-F) Representative images of specific neurite morphology defects observed in touch cells. White dashed circles denote branching in panel C, an arrow points to a misguided neurite in panel D, and white stars illustrate beads in panel E and gaps in panel F, respectively. The scale bar in panel F is 10 µm. (G-L) Quantification of the defects exemplified in panels B-F in Dendra2, TauT4, and T231A, T231E and K274/281Q. The colored bar denote the percentage of worms with the defect, while the gray bar denotes the percentage of worm that lack the defect. Statistical analysis was by Fisher’s exact test followed by two-tailed correction, with *\*P<0.05* compared to the Dendra2 control. Not all significant statistical comparisons are annotated, and data for the parental *Pmec-4::mCherry* reporter strain lacking tau transgenes, which is very similar to Dendra2, is not shown. N= 25 ± 5 neurites from separate animals scored for each type of defect.

In addition to overextension defect, other neuritic abnormalities develop with age, such as branching, guidance defects, beading, and breakage (Fig. 3C-F). While none of the strains were significantly different in terms of these defects at day 3 (data not shown), both T231E and K274/281Q exhibited an increased incidence of overextension, guidance, and gap defects at day 10, but were not different with respect to branching or beading compared to the Dendra2, TauT4 or T231A mutant strains (Fig. 3H-K). It was intriguing that age exacerbated the overlap defect in K274/281Q (*p = 0.05*, between day 3 and day 10), which mirrored its effect on touch sensitivity, but that T231E had reached its maximum penetrance by day 3 of adulthood. These results suggested to us that this model is appropriate to detect subtle differences in pathology and “disease” progression as a function of specific tau PTMs.

### Mitochondrial fragmentation in tau PTM mutants

Impaired mitochondrial dynamics and excessive fragmentation have been observed in AD postmortem brains and in AD mouse models (61–63). However, the effects of disease-relevant, site-specific phosphorylated or acetylated tau on mitochondrial morphology in neurons have not been thoroughly studied in the absence of tau overexpression. To investigate whether a causal relationship exists between tau PTMs and mitochondrial morphology, we examined the mitochondrial network in the PLM cell bodies at day 3 and at day 10 of adulthood in our tau PTM mutant models. Touch cell mitochondria were labeled with mito-mKeima, a pH-sensitive fluorescent biosensor (56, 57). Mito-mKeima can be used as a dual excitation ratiometric mitophagy reporter, as we expand upon below (Fig. 5 and 6). However, here we used single wavelength excitation-emission imaging of mito-mKeima in the appropriate channel to visualize mitochondrial structure, such as shown in Fig. 4. Under these image acquisition conditions, mitochondria are visible, but mitochondria that have been engulfed by acidic vesicles are not (for convenience, heretofore we will refer to these structures as “mitolysosomes”). Based upon these images, mitochondria were categorized into four levels, from normal tubular-reticular morphology through increasing degrees of fragmentation (Fig. 4A-D and Methods). Neuronal mitochondria from day 3 adult animals had generally tubular-reticular morphology, and their distribution was independent of tau genotype (data not shown). However, by day 10 of adulthood, all of the strains contained some fragmented mitochondria, consistent with age-associated remodeling, but it was clear that T231E and K274/281Q were significantly more fragmented than Dendra2, TauT4 or T231A (Fig. 4E).

**Figure 4.**
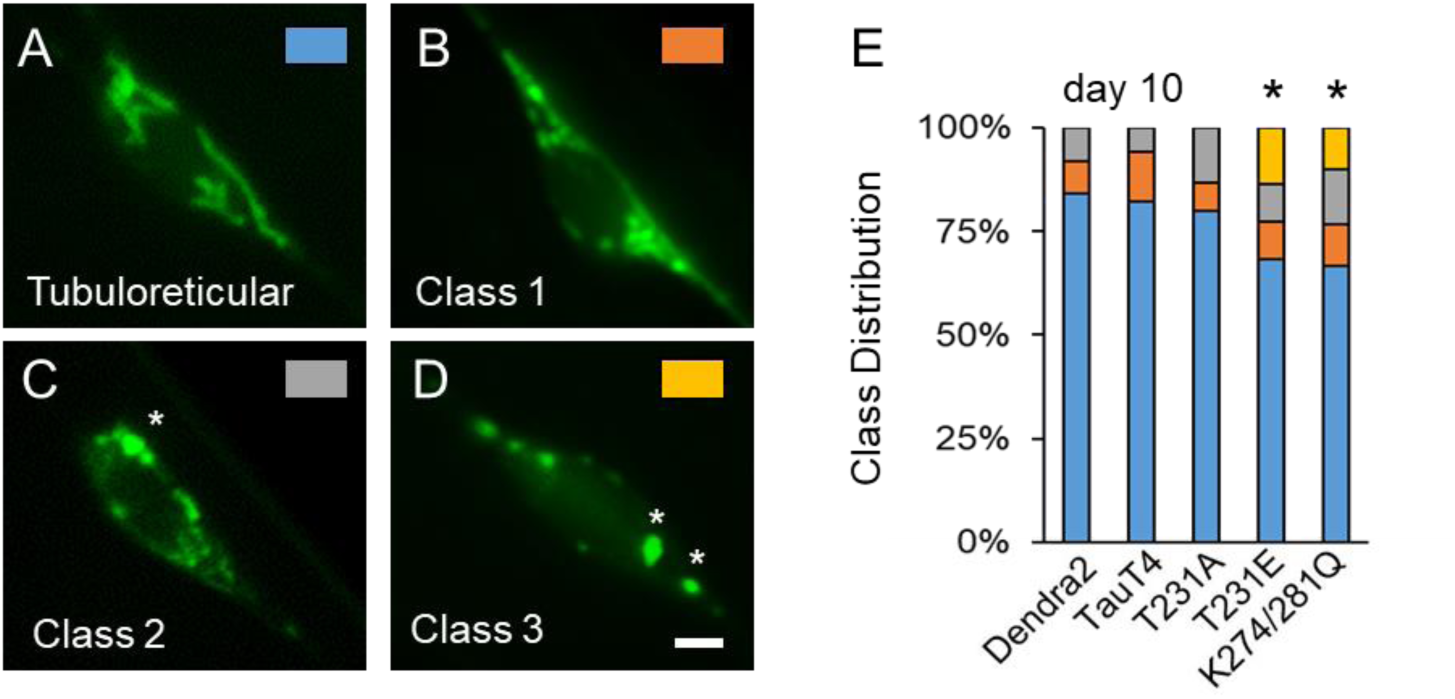
Tau PTM mutants cause mitochondrial fragmentation. (A-D) Representative images of mitochondria from PLM neuron cell bodies showing different classes of fragmentation. Each panel is color-coded to the data in panel E, as indicated. Asterisks denote overt swollen mitochondria resulting from excessive fragmentation. Labeling was via mito-mKeima, imaged on a single channel specific for mitochondria. (E) Data from day 10 adults presented in a more granular fashion, with individual cells assigned a category as depicted in panels A-D. N = 30 ± 5 cells from separate animals. Statistical analysis was by Wilcoxon signed-rank test, with *\*P<0.05* compared to Dendra2.

**Figure 5.**
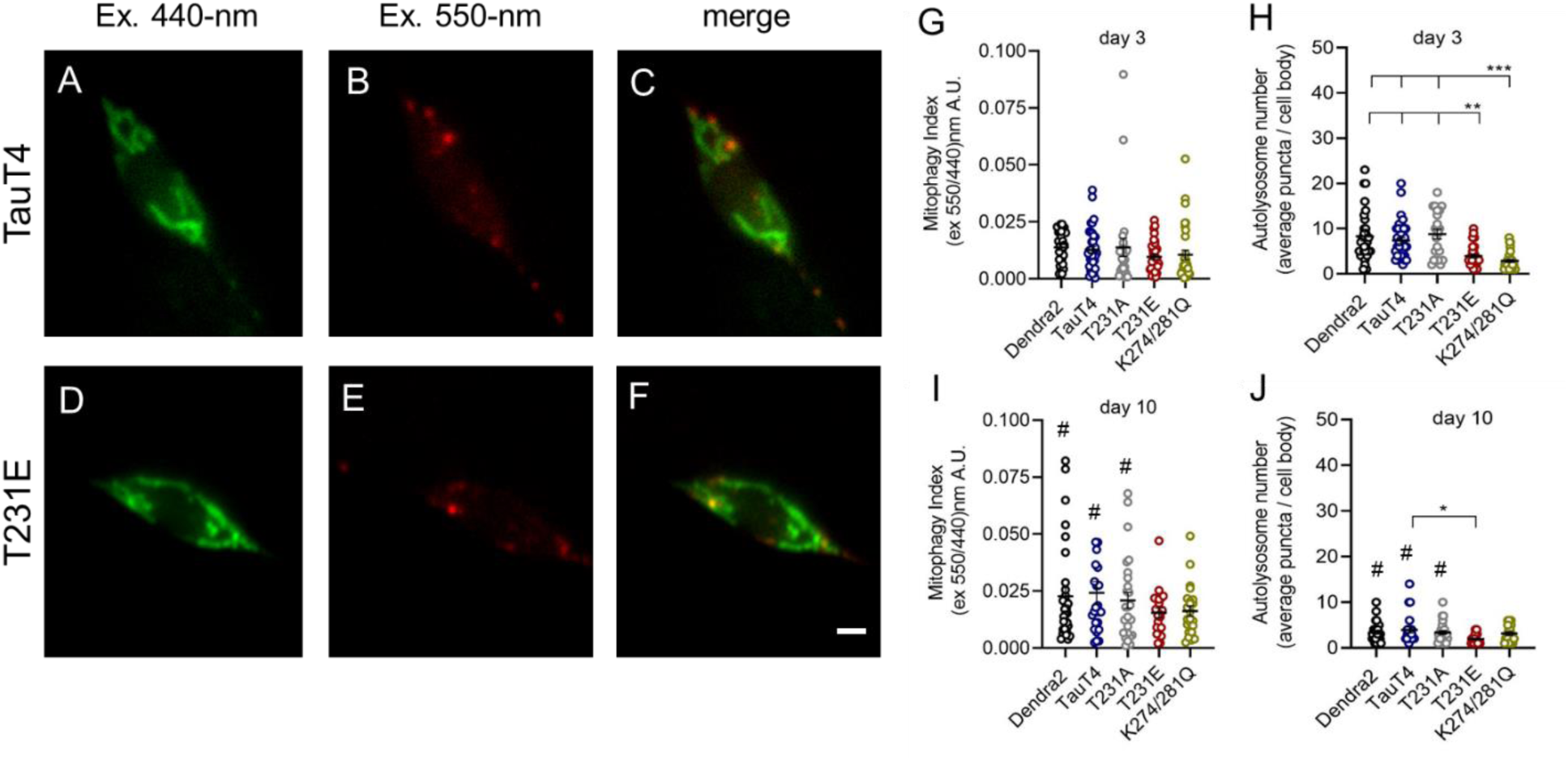
Tau PTM mutants reduce the number of mitolysosomes, but not baseline mitophagy. (A-F) Representative fluorescent images from the PLM cell bodies expressing single-copy TauT4 or T231E, together with the biosensor mito-mKeima. Mitochondria at neutral pH have been pseudo-colored green, and organelles that have been incorporated via mitophagy into acidic vesicles have been pseudo colored red. Scale bars: 5 µm. (G, I) Background corrected 550-nm excitation / 600-nm emission values were divided by 440-nm excitation / 600-nm emission values to obtain a mitophagy index for PLM cell bodies from Dendra2, TauT4, and T231A, T231E, and K274/281Q PTM mutants at day 3 and day 10 of adulthood. (H, J) Quantitative analysis of the number of mitolysosomes containing mitochondria in the distal PLM cell bodies of day 3 and day 10 adult animals as a function of tau genotype, as indicated. Data are the mean ± SEM from three independent technical replicates performed on different days. Individual data points demarcate values from single PLM cells from separate animals (N = 35 ± 5). Statistical analysis within day 3 and day 10 datasets was by one-way ANOVA with Tukey’s multiple comparison test, with *\*\*\* P< 0.001, **P<0.01, *P<0.05* when comparing bracketed samples. Comparisons between day 3 with day 10 data were limited to within a single genotype, and significance was determined by Student’s t-test, with *^#^ P<0.05*.

**Figure 6.**
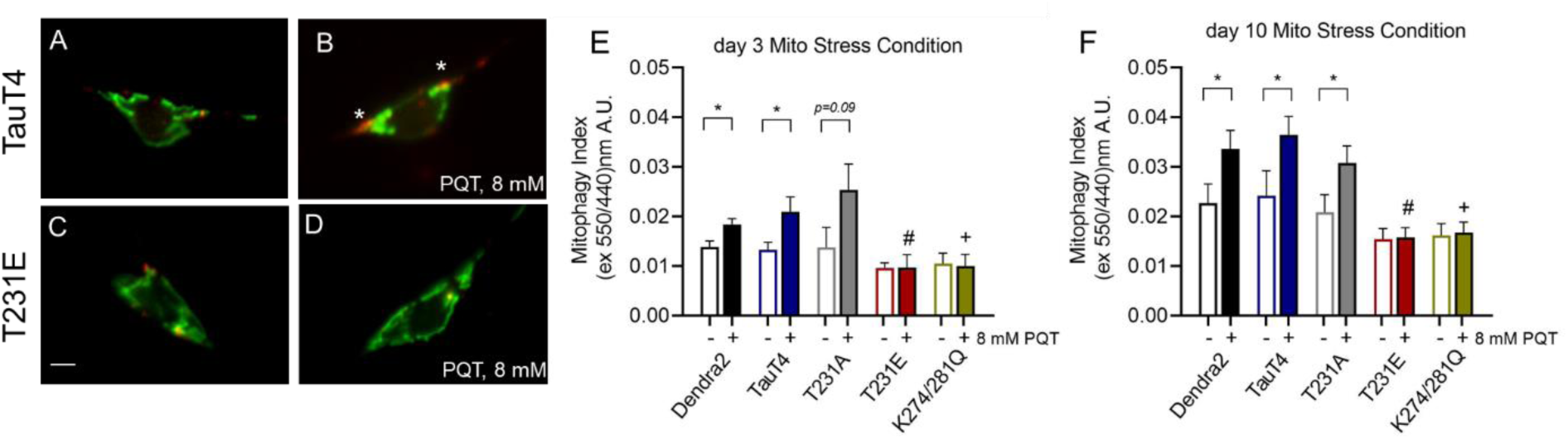
Tau PTM mutants suppress paraquat-stimulated mitophagy. mito-mKeima was used to measure mitophagy in *C. elegans* PLM touch cells following an overnight exposure to 8 mM PQT in Dendra2, TauT4, and PTM mutant strains. (A-D) are representative merged images where 440-nm excitation was used to detect mitochondria (green) and 550-nm excitation was used to detect mitolysosomes (red). Asterisks in panel B point to mitolysosomes that are clearly more abundant following PQT treatment in TauT4 animals. (E, F) Quantitative analysis of mitophagy in transgenic worms treated with PQT (8 mM overnight) immediately prior to day 3 and day 10 of adulthood. Scale bar: 5 µm. Data are the mean ± SEM from three independent technical replicates performed on different days (N = 35 ± 5 cells from separate animals). Statistical analysis was by two-way ANOVA followed by Tukey’s post hoc test, with *\*P < 0.05* denoting significance when comparing bracketed samples. #, + denotes *P < 0.05* between the PQT-treated T231E or K274/281Q and Dendra2, respectively.

### Pathologic tau modifications suppress stress-induced mitophagy

Next, we employed mito-mKeima in dual excitation mode in order to assess organelle turnover. Throughout a neuron’s lifetime, aged and damaged mitochondria undergo dynamic recycling and elimination (64). Mitophagy is a type of cargo-selective autophagy where defective mitochondria are engulfed by autophagosomes and subsequently degraded by fusion with lysosomes (Fig. S3) (65). This process of mitochondrial quality control (MQC) can be impaired during aging and has been associated with major neurodegenerative disorders including AD (66, 67). Mito-mKeima has a unique spectral characteristic whereby at neutral pH or above, such as occurs in the mitochondrial matrix, excitation at 440-nm results in emission at 600-nm+, but at acidic pH, such as occurs in the lysosome, the excitation maxima shifts to 550-nm (akin to a conventional red fluorescent protein). Mito-mKeima is also resistant to degradation by lysosomal proteases. These characteristics allows a mitophagy index to be calculated using dual excitation ratio imaging that reflects the relative amount of mitochondria that have undergone engulfment and fusion with acidic vesicles. In addition, because these mitolysosomes are spectrally and morphologically distinct (Fig. 5), we can also assess their absolute abundance and size.

In PLM neurons, pathologic tau modifications T231E and K274/281Q had little effect on baseline mitophagy, but decreased the number of mitolysosomes in young adults (Fig. 5). We also note an apparent increase in the mitophagy index and reduction in the number of mitolysosomes with age that reached significance in Dendra2, TauT4, and T231A, but not in T231E and K274/281Q (Fig. 5G, H, I, J).

Next, we sought to evaluate the impact of oxidative stress on neuronal mitophagy. These studies are particularly significant, as chronic mitochondrial stress is likely to be a factor in neurodegenerative diseases including AD (68). To induce mitophagy, Dendra 2, TauT4 and PTM mutant strains expressing mito-mKeima in touch cells were treated with 8 mM mitochondrial complex I inhibitor paraquat (PQT) overnight (Fig. S4). PQT has been used extensively in worms, including for this purpose (51, 52). The next day, mitophagy was assessed through dual-excitation ratio imaging. Unsurprisingly, PQT treatment increased mitophagy in Dendra2 at both day 3 and day 10 of adulthood (Fig. 6A, B, E, F). As found for previous measures, TauT4 and T231A were indistinguishable from Dendra2 (Fig. 6E, F). However, PQT-induced mitophagy was abolished in T231E and K274/281Q at both day 3 and day 10 (compare Fig. 6B and D as well as 6E, F). We conclude that site-specific phosphorylation and/or acetylation of tau, in addition to being a mitocentric stress in-and-of itself, has the ability to reduce normal mitochondrial responses to subsequent stress, which could impact mitochondrial function and neuronal health during aging.

## DISCUSSION

A characteristic hallmark of the AD brain is the presence of tau with PTMs defined as pathological, that likely contribute to the onset and progression of the disease. Phosphorylation of tau at specific epitopes is widely appreciated to contribute to AD (6, 8), with acetylation of tau at specific sites also shown to contribute to the evolution of tau pathology (18, 19). While it has now become evident that the insoluble accumulations of tau in the AD are likely not the primary toxic species (5–7, 69), the specific mechanisms by which monomers or soluble oligomers of tau with AD-relevant PTMs cause neuronal dysfunction have not been full delineated. This is due in part to the fact that the majority of studies have used models in which tau is overexpressed, which can result in outcomes that may not be directly relevant to AD pathogenesis. To avoid this potential confounding factor, we generated *C. elegans* strains containing single-copy expression cassettes coding for tau and tau with AD-associated PTMs. These transgenic animals allowed us to make several key discoveries, which provide important insights into the mechanisms by which tau with AD-relevant PTMs when expressed at physiological levels may impair neuron function.

Although 0N4R human brain tau isoform contains almost 70 potential phosphorylation sites that span the entire molecule (4), only select residues are phosphorylated physiologically and/or pathologically. One key disease-relevant site is T231 that shows increased phosphorylation early in the evolution of AD tau pathology and greater levels in “pre-tangle” neurons (8). Phosphorylation of T231 results in a decrease in microtubule association (70), likely due to the conformational shift and decreased tubulin binding that was observed with a pseudophosphorylated tau construct (10). A S235/T231E tau construct also showed mislocalization in mature neurons (71). Intriguingly, we observed that at day 3 worms expressing T231E showed subtle but significant defects in touch sensitivity and neurite morphology, while those expressing the acetylation mimic K274/281Q did not. However, by day 10 significant deficits in touch sensitivity and neurite morphology were observed in both T231E and K274/281Q. Thus functional decline of the touch neurons due to tau modifications is highly correlated with altered neuron morphology providing hints towards commonalities with the aging mammalian brain and suggesting conserved mechanisms can be operative in neuronal decline across phyla (72). Overall these data may suggest that phosphorylation at T231 is an early initiator of tau dysfunction in AD. These findings also correlate with the fact that increases in phosphorylation at T231 precede increased acetylation at K274/281 in the evolution of AD tau pathology (4, 73).

Mitochondria are crucial metabolic hubs dictating cell fate decisions, and mitochondrial dysfunction likely plays a critical role in the pathogenesis of AD (23, 24, 72). Mitochondria possess dedicated MQC mechanisms to ensure their fidelity (65). Abnormalities in MQC pathways noted to occur in AD (64) may arise in part through the action of tau species with aberrant PTMs (74). Mitophagy, which is a form of selective autophagy that delivers dysfunctional mitochondria to lysosomes for recycling, is a key player in MQC. *C. elegans* have been widely used to study neuronal function, aging, and MQC mechanisms, as well as to model proteotoxic neurodegenerative disorders (75). Therefore, we next examined the impact of T231E and K274/281Q on mitochondrial biology. In contrast to the deficits in touch sensitivity and neuronal morphology observed at day 3, neither T231E nor K274/281Q negatively impacted mitochondrial morphology at that age. However, by day 10 mitochondrial fragmentation was significantly exacerbated in T231E and K274/281Q, which could be reflective of a deficit in mitophagy. Therefore, we measured the relative amount of mitochondria that were engulfed and fused with acidic compartments, as well as the absolute abundance of mitochondria in acidic compartments (“mitolysosomes”).

Interestingly, we found that in control animals, the mitophagy index, a measure of relative mitolysosome to mitochondria abundance, increased with age, and the number of mitolysosomes decreased (Fig. 5). The factors critical for the effective turnover of damaged mitochondria during aging likely include underlying stress, as well as autophagic and lysosomal capacities. While there are other reports of mitophagy increasing with age in systems including Drosophila (76), mouse (77) and human disease (78), there is also evidence that mitophagy becomes, like many types of stress responses, impaired with age (79). Although our data support the former studies, we note that mito-mKeima is resistant to acid proteases and likely accumulates over time. In fact, a decreased number of brighter mitolysosomes in day 10 animals may represent cumulative vesicle fusion, and so we need to temper our conclusion to reflect this caveat. Nevertheless, we were able to stimulate mitophagy using PQT at day 10 to a similar extent as day 3 (Fig. 6), confirming that, at a minimum, the ability to generate a robust response to oxidative stress is maintained in older wild type animals.

Our results demonstrate a striking abolition of PQT-induced mitophagy in the AD-relevant T231E and K274/281Q mutants (Fig. 6). This observation is consistent with defective mitophagy being a prominent feature in age-related disorders (80), including AD (81), and contributing to premature aging such as observed in Werner’s syndrome patients and invertebrate Werner’s disease models (82). It is also particularly intriguing that the T231E and K274/281Q do not appear to exhibit the same age-dependence as Dendra2, TauT4, or T231A. This could be interpreted to mean that these mutants exhibit characteristics that appear in older adults. Their inability to response appropriately to oxidative stress – at both a young and old age - suggests that the mitochondria in fact do have baseline defects, albeit at a level that is not discernable in the absence of stress. The recent finding that mitophagy enhancement can suppress AD-related phenotypes in tau transgenic animals lends support to this idea (83).

It will be of interest to determine whether the tau mutants described here are perceived as stressors, and hence cause activation of a retrograde response, such as has been described previously for the *C. elegans* Nrf2 ortholog SKN-1 in adaptation to a decrease in mitophagy (84). Alternatively, other retrograde signaling pathways such as that mediated by ATFS-1 and the mitochondrial unfolded protein response (mtUPR) can also mediate adaptation to mitochondrial stress (85), including stress due to defects in mitophagy machinery (86). However, prolonged cellular activation of the mtUPR has been shown to be maladaptive in a *C. elegans* model of dopaminergic neurodegeneration (87), suggesting that ultimate role of stress response pathways is context dependent. It is also possible that the single-copy tau mutants do not elicit stress-responses in-and-of themselves, but instead sensitize neurons to additional stressors, consistent with our mitophagy results.

## CONCLUSION

In conclusion, to our knowledge this is the first study to clearly demonstrate that single copy expression of tau with AD-associated PTMs impairs neuronal function and structure in an age-dependent manner. In addition, the effect of tau modifications on stress-induced mitophagy could lead to cumulative metabolic defects and energetic crises with age. One advantage of our single-copy model is that it allows us to quantitatively measure subtle deficits and discriminate between the effects of distinct PTMs. For example, we demonstrate that T231E presents with a neuronal functional (and morphological) deficit earlier than K274/281Q. Since stress-induced mitophagy was abolished equally by both, it is likely that AD-associated, pathologic tau mutants are differentially impacting neuron structure/function through at least one other mechanism. However, further studies are needed to determine if these pathways are separate and isolated, and if they interact, whether they are additive or synergistic. We anticipate that this new *C. elegans* AD model represents a foundation to achieve a more nuanced understanding of how tau PTMs impact neuronal function.

## Supporting information

Four figures and two tables

## Abbreviations

AD: Alzheimer’s disease
CRISPR: clustered regularly interspaced short palindromic repeats
Cas9: CRISPR associated protein 9
ETC: electron transport chain
MQC: Mitochondrial quality control
MosSCI: Mos-mediated single copy insertion
NGM: nematode growth media
PQT: paraquat
PTM: post translational modification

## DECLARATIONS

### Availability of Data and Materials

The datasets used and/or analysed during the current study are available from the corresponding author on reasonable request.

## Author contributions

Conceived and designed the experiments: SG GJ KN. Performed the experiments: SG SF. Analyzed the data: SG SF GJ KN. Wrote the paper: SG GJ KN. All authors read and approved the final manuscript

## Acknowledgements

Technical assistance provided by Teresa Sherman, Joseph Cartella, Alan Alberto and Ejiroghene Okarevu was greatly appreciated. We thank the members of Dr. Johnson’s lab, the Mitochondrial Research and Interest Group at the University of Rochester Medical Center and the members of the Western New York worm meeting for their valuable suggestions and helpful discussions.

