## Supplementary material for "Alzheimer’s disease-relevant tau modifications selectively impact neurodegeneration and mitophagy in a novel *C. elegans* single-copy transgenic model": Four figures and two tables

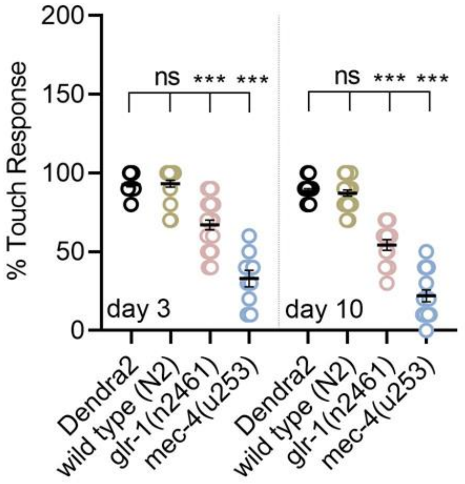


**Figure S1. Single-copy Dendra2 transgenic worms are indistinguishable from the N2-Bristol wild-type control strain with respect to touch sensitivity.** Touch assays were conducted on day 3 and day 10 adult animals grown at $20^0$C using *glr-1(n2461)* and *mec-4(u253)* strains as negative controls. The data are the mean ± SEM (N = 10-25 animals). Statistical analysis was by one-way ANOVA followed by Tukey's multiple-comparisons test with **P < 0.001* denoting significance between bracketed samples (ns is not significant). Each point represents a value obtained from a single animal.

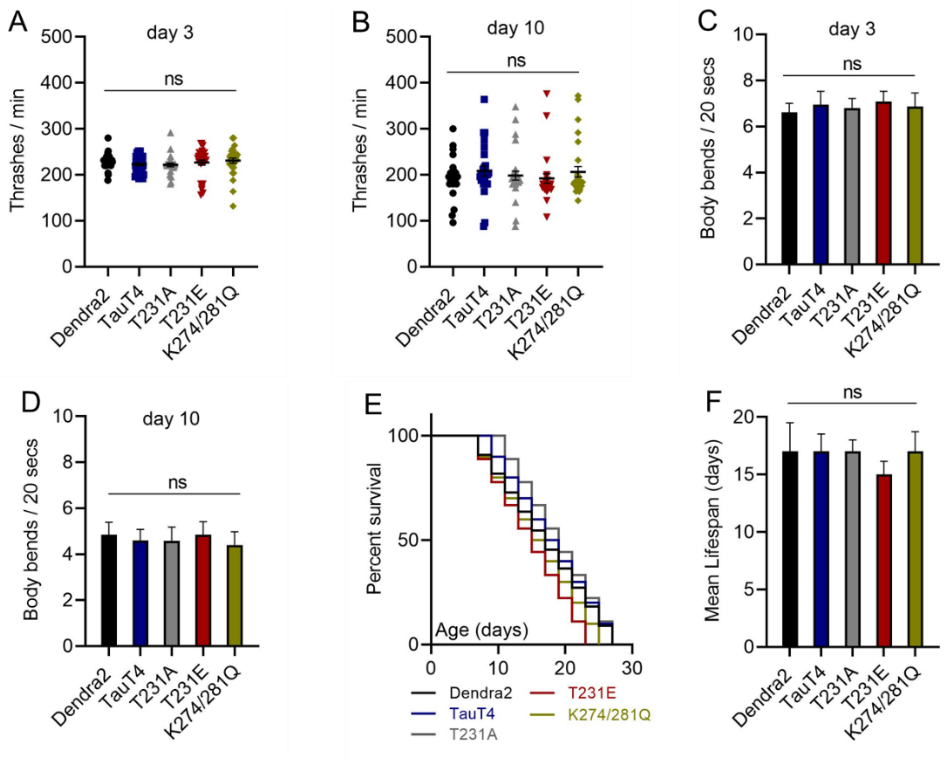


**Figure S2. Thrashing, locomotion, and lifespan**. Dendra2, TauT4, T231A, T231E and K274/281Q PTM mutant strains at day 3 and at day 10 of adulthood were assessed for (A, B) thrashing behavior in liquid media, (C, D) locomotion on solid media measured as the rate of body bends, (E) survival, or (F) lifespan. The data are the mean ± SEM. (N=20-to-50 worms per genotype, with each data point in A, B representing an individual worm). Statistical analyses were by one-way ANOVA or Mantel-Cox/ log-rank analysis. No statistical differences were found between genotypes for any of the three measures (denoted ns).

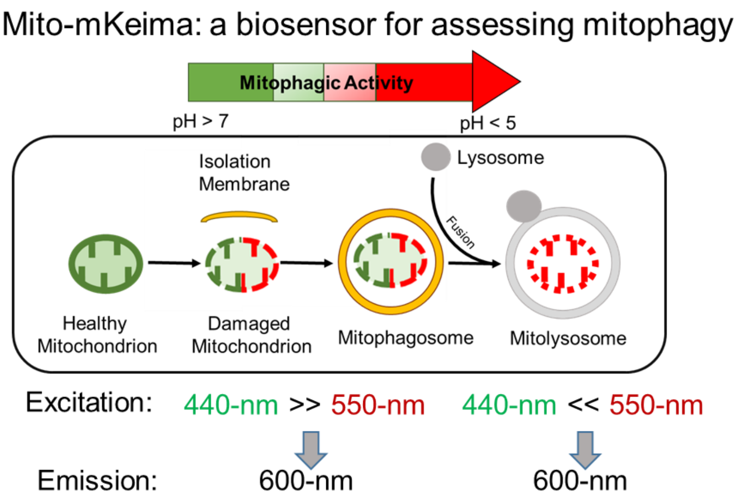


**Figure S3. Schematic representation of using mito-mKeima to measure mitophagy.** Healthy mitochondria have a matrix pH > 7 (green) whereas dysfunctional mitochondria undergoing mitophagy and engulfed by lysosome are exposed to an acidic pH < 5 (red). Mito-mKeima is an acid-protease resistant fluorescent protein whose excitation maxima shifts from 440-nm to 550-nm with acidification. By measuring the ratio of emissions at 600-nm following sequential excitations at the two excitation maxima, you can obtain an estimate of the relative amount of mitochondria that are in neutral compared to acidic environments. In addition to spectral differences, mitochondria that have been engulfed by autophagosomes (labeled mitolysosomes) are generally round rather than tubuloreticular, and hence morphology can generally be used as a second measure to distinguish between organelles. However, this difference can be masked by mitochondrial fragmentation.

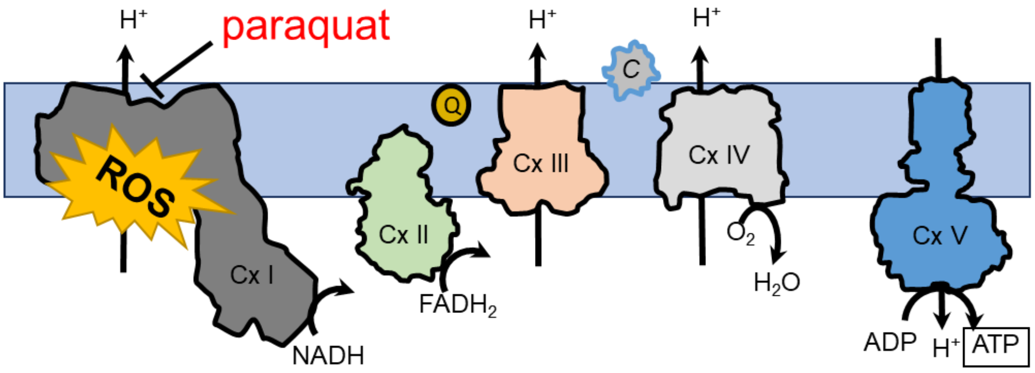


**Figure S4. Schematic of the electron transport chain (ETC).** The ETC consists of complexes I-V, with substrate entry occurring at Cx I and Cx II, proton pumping occurring at Cx I, Cx III, and Cx IV, and ATP synthesis occurring at Cx V. Coenzyme Q (Q) and cytochrome C (C) are shown as well. Targeting of Cx I by paraquat generates reactive oxygen species (ROS) and results in oxidative stress.

| Strain | Transgene | Description |
| --- | --- | --- |
| KWN169 | *(Pmec-7::Dendra2)* | No tau |
| KWN167 | *(Pmec-7::Dendra2::TauT4)* | Wild type tau (0N4R) |
| KWN788 | *(Pmec-7::Dendra2::TauT4(T231A))* | Phospho-ablation |
| KWN789 | *(Pmec-7::Dendra2::TauT4(T231E))* | Phospho-mimetic |
| KWN790 | *(Pmec-7::Dendra2::TauT4(K274/281Q))* | Acetyl-mimetic |

**Table S1: Tau PTM mutant transgenes.** All tau transgenes are translational fusion to photo-convertible protein Dendra2. 0N4R is the wild type human tau isoform used in this study. CRISPR-Cas9 gene editing was used to introduce phosphomimetic T231E, phosphoablation T231A, and acetylmimetic K274/281Q mutations into the TauT4 ORF. For simplicity, these mutants will be referred to as T231E, T231A and K274/281Q.

| Transgene | Penetrance of defects in day 10 adults | | | | |
|---|---|---|---|---|---|
| | *Overextended* 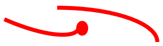 | *Branched* 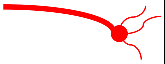 | *Misguided* 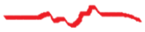 | *Beaded* 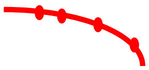 | *Gap* 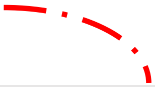 |
| Dendra2 | 2 of 20 | 1 of 20 | 2 of 20 | 2 of 20 | 1 of 20 |
| TauT4 | 3 of 20 | 1 of 20 | 2 of 20 | 3 of 20 | 3 of 20 |
| T231A | 2 of 22 | 1 of 22 | 2 of 22 | 4 of 22 | 2 of 22 |
| T231E | 13 of 30 | 3 of 30 | 11 of 30 | 6 of 30 | 13 of 30 |
| K274/281Q | 11 of 25 | 3 of 25 | 4 of 25 | 6 of 25 | 11 of 25 |

**Table S2: Detailed quantification of the neuronal defects observed in day 10 transgenic worms.** The most significant defects observed in tau PTM-mimetics are the overextension of ALM/PLM and gaps or breaks observed in their neuronal processes.
